# Cyclin A2/Cdk1 is Essential for the *in vivo* S Phase Entry by Phosphorylating Top2a

**DOI:** 10.1101/513861

**Authors:** Miaomiao Jin, Ruikun Hu, Baijie Xu, Weilai Huang, Hong Wang, Bo Chen, Jie He, Ying Cao

**Affiliations:** Clinical and Translational Research Center of Shanghai First Maternity and Infant Hospital, School of Life Sciences and Technology, Tongji University, Shanghai 200092, China.; Institute of Neuroscience, State Key Laboratory of Neuroscience, CAS Center for Excellence in Brain Science and Intelligence Technology, Chinese Academy of Sciences, Shanghai 200031, China.; University of Chinese Academy of Sciences, Beijing, China.; The MOE Key Laboratory of Cell Proliferation and Differentiation and the State Key Laboratory of Membrane Biology, College of Life Sciences, Peking University, Beijing 100081, China.

**Keywords:** Cdk1, Cyclin A2, Cyclin B1, Top2a, M phase, S phase entry, IKNM

## Abstract

Cyclin-dependent kinase 1 (CDK1) plays essential roles in cell cycle regulation. However, due to the early embryonic lethality of mouse *Cdk1* mutants, the *in vivo* role of CDK1 in regulating cell cycle and embryonic development remains unclear. Here, by generating zebrafish *cdk1* mutants using CRISPR/Cas9 system, we show that *cdk1*^−/−^ embryos exhibit severe microphthalmia accompanied with multiple defects in polarized cell division, S phase entry and M phase progression, cell apoptosis and cell differentiation, but not in interkinetic nuclear migration (IKNM). By informatics analysis, we identified Top2a as a potential downstream target, and Cyclin A2 and Cyclin B1 as partners of Cdk1 in cell cycle. Depletion of either Cyclin A2 or Top2a leads to decreased S phase entry and increased DNA damage response in zebrafish retinal cells, and depletion of Cyclin B1 leads to M phase arrest. Immunoprecipitation shows that Cdk1 and Cyclin A2 physically interact *in vivo*. Moreover, phosphorylation of Top2a on Serine 1213 (S1213) site is almost absent in either *cdk1* or *ccna2* mutants, but in not *ccnb1* mutants. Furthermore, overexpression of TOP2A^S1213^, the phosphomimetic form of human TOP2A, rescues S phase entry and microphthalmia defects in *cdk1*^−/−^ and *ccna2*^−/−^ embryos. Taken together, our data suggests that Cdk1 interacts with Cyclin A2 to regulate S phase entry through phosphorylating Top2a, and with Cyclin B1 to regulate M phase progression *in vivo*.

## Introduction

Cell cycle progression is tightly regulated by cyclin-dependent kinases (CDKs) (Malumbres, 2014; Malumbres and Barbacid, 2005). Extensive studies have identified more than 20 CDKs, which can be further categorized as interphase CDKs and mitotic CDKs (Malumbres, 2014). However, genetic evidences revealed different roles for CDKs in animal development and cell cycle regulation. Depletion of an interphase CDK, including CDK2, 3, 4, 6, does not affect mice survival (Berthet et al., 2003; Malumbres et al., 2004; Ortega et al., 2003; Ye et al., 2001). When all interphase CDKs are depleted simultaneously, the mouse embryos can still undergo organogenesis until midgestation(Santamaria et al., 2007). On the contrary, in the absence of CDK1, the mitotic CDK, the mouse embryos fail to develop to the morula and blastocyst stages(Santamaria et al., 2007), suggesting that CDK1 is the most important CDK during cell proliferation and early development. Previous findings suggested that CDK1 can compensate the roles of other CDKs during the S phase entry or G1/S phase transition in the absence of all interphase CDKs(Hochegger et al., 2007; Santamaria et al., 2007). Furthermore, activated by Cyclin E or Cyclin A2, CDK1 itself is required for the S phase entry in mouse fibroblast cells by regulating the firing program of DNA replications (Katsuno et al., 2009; Nakanishi et al., 2010). However, the *in vivo* function and mechanism of CDK1 in cell cycle regulation is still lacking.

Here, we took advantage of the zebrafish model to address this question by characterizing Cdk1-depleted embryos. We found that Cdk1 is required for retinal development by regulating S phase entry and M phase progression. By analyzing the database of Cdk1 substrates and genes essential for retinal development, we identified Top2a as a potential downstream factor of Cdk1. *Top2a*^−/−^ mutants phenocopy *cdk1*^−/−^ mutants and the phosphorylation level of Top2a is reduced in *cdk1*^−/−^ mutants. Importantly, when overexpressed with Top2a^S1213D^, a mutant form of Top2a with S1213 replaced by a phosphomimetic residue (S1213D), can partially rescue the microphthalmia and S phase entry defects of *cdk1*^−/−^ mutants. These results suggested that Top2a functions downstream of Cdk1 in S phase entry. As both Cyclin A2 and Cyclin B1 are required for retinal development, we asked what the in *vivo* functions of these two Cyclins are during cell cycle. We discovered that Cyclin A2 physically interacts with Cdk1, and is also required for S phase entry, but not M phase progression. Cyclin A2-depleted embryos phenocopy *cdk1*^−/−^ mutants, and can also be rescued by Top2a^S1213D^ overexpression, which suggested that Cdk1 functions together with Cyclin A2 in regulating S phase entry through Top2a. In the contrary, Cyclin B1 is required for M phase progression. Taken together, Cdk1 interacts with different Cyclins in regulating the *in vivo* progression of S phase and M phase.

## Results

### Knockout of *cdk1* in zebrafish leads to degenerated retina

To study the function of Cdk1, we took advantage of the CRISPR/Cas9 system(Ota et al., 2014) to generate *cdk1* knock-out mutants in zebrafish. The DNA sequence of the mutated *cdk1* allele carries a premature stop that results in a truncated form of Cdk1 protein (Fig. 1A). In contrast to *cdk1*^−/−^ mice, which die before blastula stage, zebrafish *cdk1*^−/−^ embryos survived until 5 days post fertilization (dpf), which is likely due to the maternal deposit of Cdk1 mRNA (Supplementary Fig. 1A). Western blot showed that Cdk1 protein was completely abolished in homozygotic *cdk1*^−/−^ at 24 hours post-fertilization (hpf), the time at which the mutant embryos were morphologically indistinguishable from the wild-type embryos (Fig. 1B). The *cdk1^−/−^*mutants started to show microphthalmia at about 30 hpf and progressed to severely degenerated retina at 72 hpf (Fig. 1C). The microphthalmia phenotype was segregated with the mutant *cdk1* allele and can be rescued by overexpressing *cdk1-gfp* mRNA (Fig. 1C), indicating that this phenotype in *cdk1*^−/−^ is specifically caused by the depletion of Cdk1. The rescue effect of Cdk1 overexpression in *cdk1*^−/−^ was further confirmed by expression of *mz98*, a retina peripheral marker(Burrows et al., 2015). The retinal expression of *mz98*, which was absent in *cdk1*^−/−^ mutants, was restored in Cdk1-GFP overexpressed *cdk1*^−/−^ embryos (Fig. 1D).

**Figure 1.**
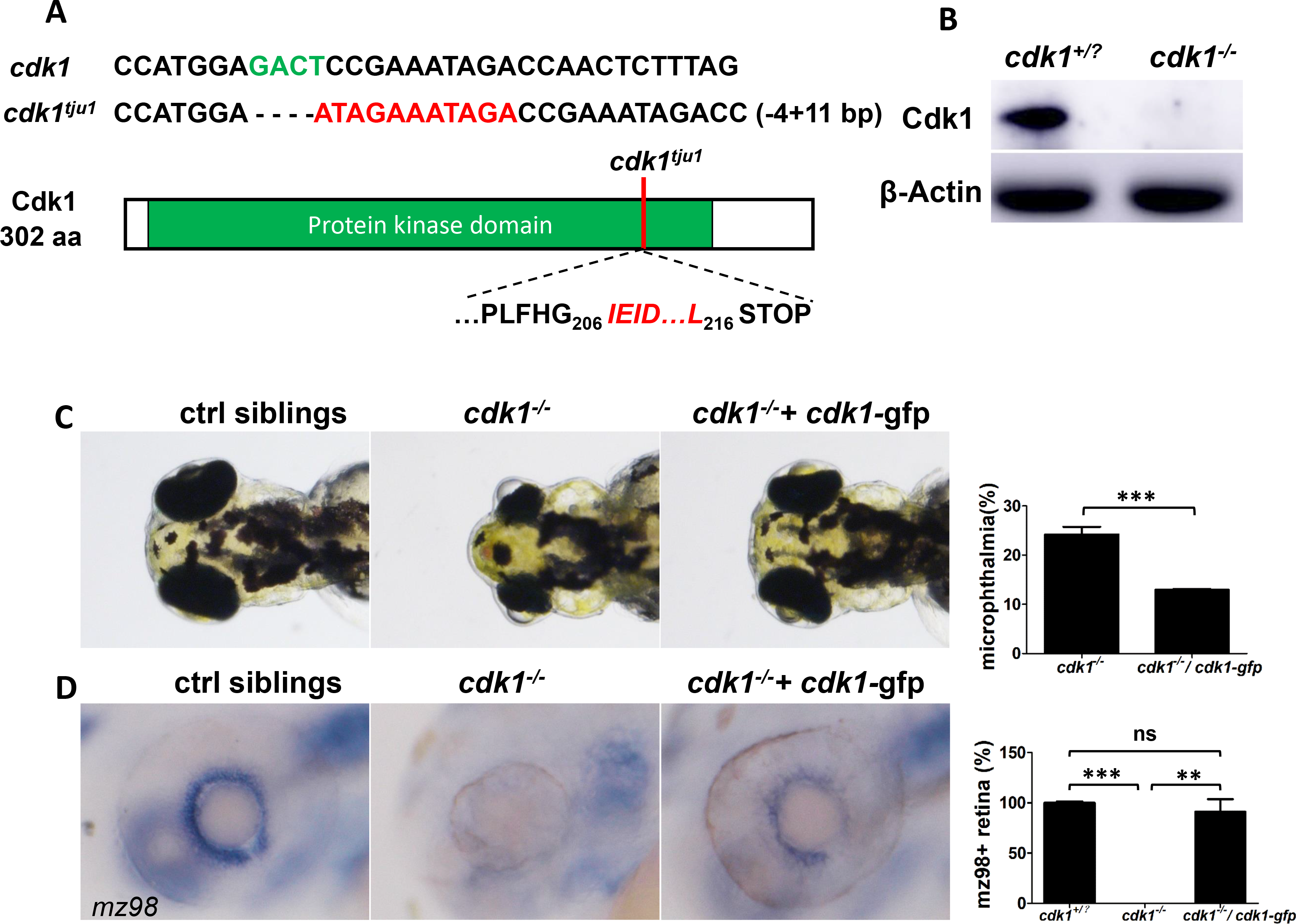
*cdk1*^−/−^ zebrafish mutants show microphthalmia. (A) Sequence information of wild-type and mutant *cdk1* alleles. Schematic of Cdk1 protein indicates position of the *cdk1*^*tju1*^ frameshift. (B) Western blot shows that Cdk1 protein is lost in *cdk1*^*−/−*^ embryos at 24 hpf. (C) Representative images of wild-type, *cdk1*^−/−^ and *cdk1*^−/−^ injected with *cdk1-gfp* mRNA embryos shows the microphthalmia phenotype of *cdk1*^−/−^ can be rescued by overexpression of *cdk1-gfp* mRNA. The percentage of embryos with microphthalmia was quantified (n≥8) and plotted. (D) Representative images of *in situ* hybridization using probe against *mz98*, a marker gene for retinal stem cells, of wild-type, *cdk1*^−/−^ and *cdk1*^−/−^ injected with *cdk1-gfp* mRNA embryos. Note that defects of retinal stem cells in *cdk1*^−/−^ can be rescued by overexpression of *cdk1-gfp* mRNA. The percentage of embryos with *mz98* positive staining in retina was quantified (n≥1) and plotted. Error bars represent S.D.; \**P*<0.05, \*\**P*<0.01, \*\*\**P*<0.001. N indicates the number of independent experiments and n indicates sample size in each group, usually the number of embryos used, unless otherwise stated.

### Cdk1 is required for cell fate differentiation in the zebrafish retina

To figure out the cellular mechanism underlying retinal degeneration of *cdk1^−/−^*, we further characterized this mutant in detail. The zebrafish retinas start to laminate at 48 hpf (Schmitt. and Dowling., 1999). Histological analysis by hematoxylin staining revealed disrupted lamination and nuclei condensation in *cdk1*^−/−^ fish retina, which suggested apoptotic cells (Supplementary Fig. 2A). We performed TUNEL assay to measure the apoptosis level in the retina (Gorczyca et al., 1993; Lozano. et al., 2009). The number of apoptotic cells labeled by TUNEL staining was significantly increased in the mutant retinas, compared with that in the control siblings at 30 hpf and 72hpf, but not at 24 hpf (Supplementary Fig. 2B). This result suggested that Cdk1 is essential for retinal development.

We further analyzed if the retinal cells can differentiate properly. Zebrafish retinas are composed of five groups of neuron cells, including photoreceptor cells (PR), bipolar cells (BCs), retinal ganglion cells (RGCs), Amacrine cells (ACs) and horizontal cells (HCs)(Schmitt. and Dowling., 1999; Zagozewski et al., 2014). Compared with the wild type retinas, the photoreceptor cells, labeled by Zpr1 or Zpr3 antibodies, were almost disappeared in the *cdk1*^−/−^ retinas (Supplementary Fig. 2C). We also took advantage of a transgenic fish *Tg*(*ath5*: *gapRFP;ptf1α: cytGFP)*, in which the cell membrane of RGCs is labeled with RFP, driven by a RGC-specific *ath5* promotor, and the cytoplasm of ACs and HCs is labeled with GFP, driven by AC- and HC-specific *ptf1a* promoter(Almeida. et al., 2014). Compared with control siblings, both RFP- or GFP-positive cells were significantly reduced from the very beginning in *cdk1*^−/−^ retinas (Supplementary Fig. 2D), suggesting that RGCs, ACs and HCs all failed to differentiate in *cdk1*^−/−^. Taken together, the differentiation of retinal neurons is disrupted in Cdk1-depleted retina.

### Cdk1 depletion leads to oriented cell division defects in the zebrafish retina

As Cdk1 is a cell cycle regulator, we examined the cell proliferation status of the retina in the *cdk1*^−/−^ mutants by immunostaining with an antibody against Phosphor-Histone H3 (pH3), which labels the mitotic cells at M phase. Surprisingly, the mitotic cell number in the *cdk1*^−/−^ retina was comparable to that in the control siblings at 24 hpf and 30 hpf (Fig. 2, A - D). Nevertheless, the distribution pattern of mitotic cells was different between the retinas of the *cdk1*^−/−^ and control embryos at 30 hpf. The dividing cells were only observed at the apical side in control siblings, but in the *cdk1*^−/−^ mutants, dividing cells were also aberrantly localized to the basal of the retina (Fig. 2C).

**Figure 2.**
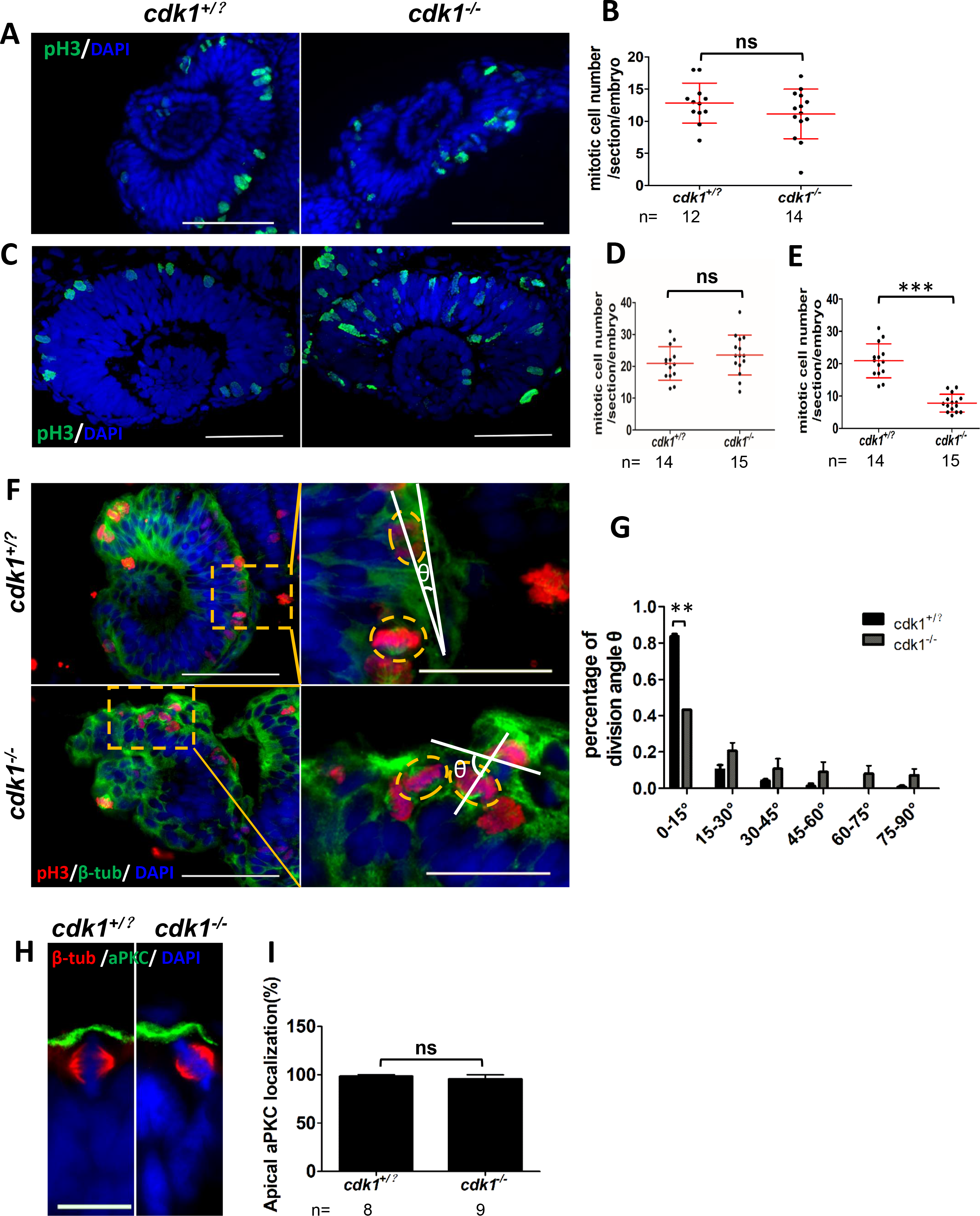
Cell division defects in *cdk1*^−/−^ retina. (A, C, F, H) Embryos were sectioned, and subjected to IF staining with indicated antibodies. (A, B) Mitotic cells (pH3) remained unchanged in *cdk1^−/−^*embryos at 24 hpf (n≥5 embryos). (C) The distribution pattern of mitotic cells (pH3) was shifted from the apical to the basal in *cdk1*^−/−^ retinas at 30 hpf. Scale bars, 50 μm. (D, E) pH3 positive cell number in the whole retina (D) or at the apical of the retina (E) per section in each embryo was counted and plotted (n>15 sections from 5-8 embryos). (F) Oriented cell division was disrupted in *cdk1*^−/−^ retinas at 24 hpf. Right panels show magnification of the yellow boxed region in the left panels. The division angle θdefined as the angle between the cell division axis and the tangent line of the retina. (G) The division angles were measured and plotted (≥90 angles from 16 embryos). Scale bars, 50 μm in the left and 20 μm in the right. (H) aPKC localization in dividing cells (β-Tub) remained unchanged in *cdk1*^−/−^ retinas at 24 hpf. Scale bars, 10 μm. (I) Percentage of normal localization of aPKC were counted and plotted. Error bars represent S.D.; ns, not significant, \*\**P*<0.01.

During zebrafish development, retinal cells divide parallelly to the tangent plane of the apical surface or horizontally (Agathocleous and Harris, 2009; Das et al. 2003; Poggi et al., 2005). We noticed that the division angles became randomized in *cdk1*^−/−^ retinas compared with controls (Fig. 2F). We further analyzed the mitotic spindle orientation by staining the chromosomes (DAPI) and spindle fibers (β-tubulin). We defined angle between the cell division axis and the tangent line of the retina as division angle. In control embryos, more than 80% of mitotic cells divided horizontally (division angle ≤ 15°), whereas only 40% cells divided horizontally in *cdk1* mutants (Fig. 2G). These results indicate that Cdk1 is required for oriented cell division.

Previous studies show that atypical PKC (aPKC) is required for proper spindle alignment (Guilgur et al., 2012; Hao et al., 2010; Vorhagen and Niessen, 2014). In the zebrafish retinas, aPKC localizes to the apical side of dividing progenitor cells and depletion of aPKC leads to randomized cell division and reduced eye size (Horne-Badovinac et al., 2001). We further investigated the subcellular location of aPKC in the *cdk1*^−/−^ retina. Compared with the wild type, the localization of aPKC remained same in *cdk1*^−/−^ mutant retinas (Fig. 2H), suggesting that the function of Cdk1 in regulating oriented cell division is aPKC-independent.

### IKNM is normal in *cdk1* mutants

During retinal development, the basally localized nuclei migrate to the apical side to divide during G2 phase, and the nuclei of daughter cells will move basally again after cell division. This process is called the interkinetic nuclear migration (IKNM) (Baye and Link, 2007, 2008; Norden et al., 2009). Since in *cdk1*^−/−^ mutants, the mitotic cells were not only localized to the apical side of the retinas, but were also aberrantly localized to the basal side (Fig. 2C), we wondered whether the IKNM process was disrupted in the mutants. To address this question, we examined the process of IKNM by live imaging in mosaically GFP-labeled *cdk1*^−/−^ embryos which were injected with *GFP* mRNA into one cell at the 16-cell stage. Noticeably, we observed that the cells rounded up at the apical surface and then divided into two cells in the control retina. We also observed the retinal cells rounded up only at the apical but not basal in the *cdk1*^−/−^ retinas, suggesting the IKNM was normal. We noticed that the number of round cells was significantly decreased in the *cdk1^−/−^* retinas compared with the control (Fig. 3A). Furthermore, the apical round cells always divided into two cells in the wild-type retinas, but in the *cdk1*^−/−^ retina, some round cells elongated and migrated basally without division (Fig. 3A). This phenomenon suggested that the basally mislocalized cells by pH3 staining (Fig. 2C) were not cells in normal division, but were cells that failed to undergo normal cytokinesis and arrested in G2 or M phase. To further confirm this observation, we immuno-stained these mosaically GFP-labeled embryos with pH3 antibody. The results showed that in *cdk1*^−/−^ retina, all apical pH3-positive cells were round cells similar to the dividing cells observed in the control retina, but the basal pH3-positive cells were elongated cells, as shown in the live imaging (Fig. 3B). These results indicated that only the apical pH3-positive cells were truly dividing cells and the basal pH3-postive cells were G2 or M phase-arrested cells that could not exit cell cycle. If we only count apical pH3-positive cells as mitotic cells, the number of mitotic cells in the *cdk1*^−/−^ retinas was significantly decreased than that in the wildtype retinas (Fig. 2 E). Taken together, in the *cdk1*^−/−^ retina, cell proliferation was significantly decreased, and basally-localized pH3-positive cells were cells caused by G2 or M phase arrest but not by aberrant IKNM.

**Figure 3.**
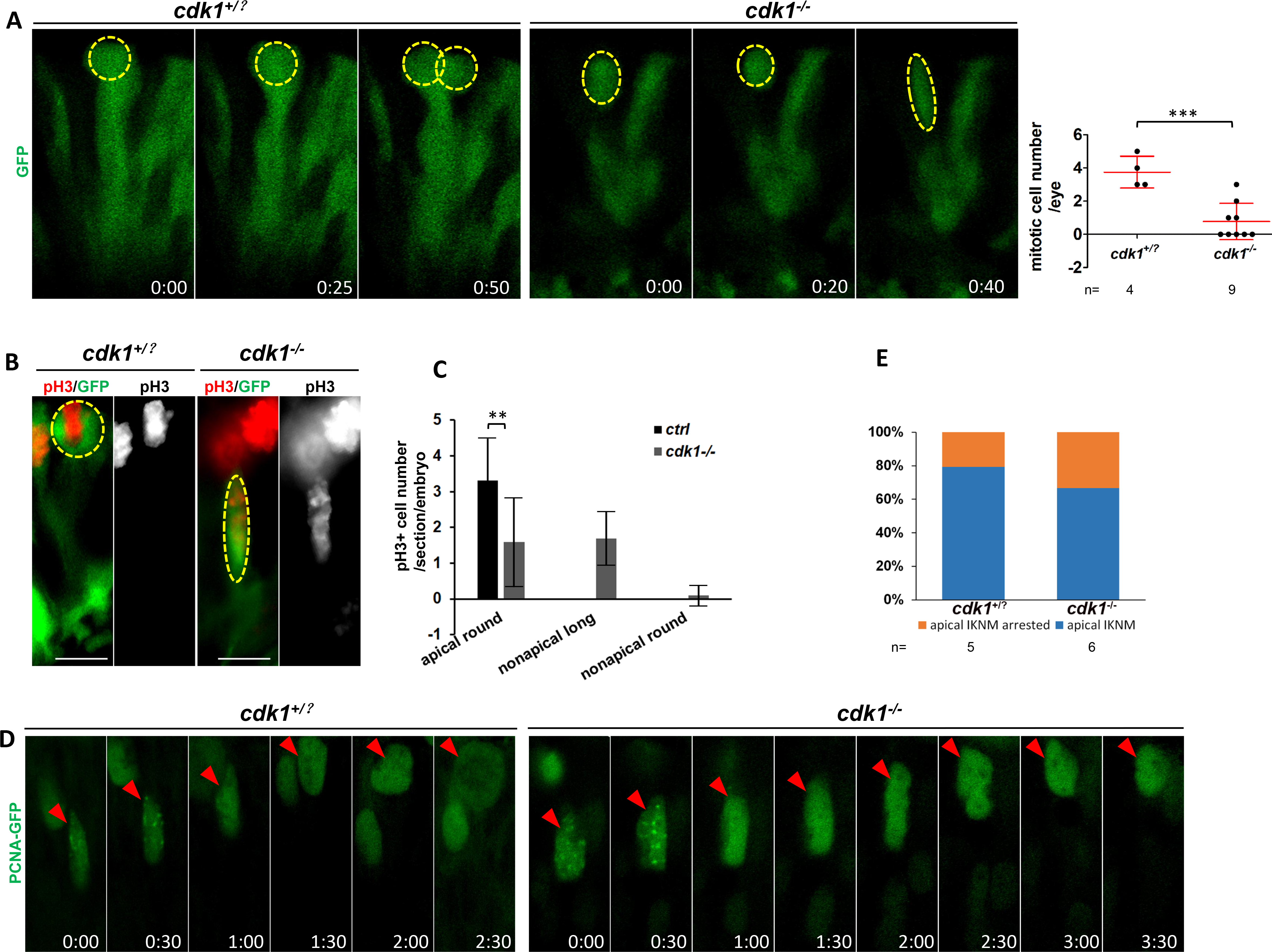
IKNM is normal in the *cdk1*^−/−^ retina. (A, B, D) Embryo retinal cells were mosaically labeled with GFP or PCNA-GFP and live-imaged (A, D), or subjected to IF staining (B). (A) GFP-labeled mitotic cells turned round at the apical (outlined by yellow dotted circles) and divided into two cells in the control sibling retinas, but in the *cdk1*^−/−^ retina some mitotic cells elongated and migrated to the basal without division. Proliferating cells (round cells at the apical) were counted and plotted according to the time-course images (n=4-9 embryos). (B) pH3+ cells were round cells at the apical in the wild-type retinas, but some pH3+ were elongated cells at the basal in the *cdk1*^−/−^ retinas, suggesting that pH3 labeled mitotic arrested cells in the *cdk1^−/−^* retinas (n≥10 eyes). Scale bars, 10 μm. (C) pH3+ cells are counted and grouped according to the shape and localization. (D) PCNA-GFP-labeled cells formed puncta and migrated apically, suggesting normal IKNM in both of the wild-type and *cdk1*^−/−^ retinas. (E) Percentage of cells with apical IKNM in wild-type and *cdk1*^−/−^ retina was counted and plotted (n=5~6 eyes). Error bars represent S.D.; \**P*<0.05, \*\**P*<0.01, \*\*\**P*<0.001.

To better observe IKNM, we mosaically labelled nuclei by overexpressing PCNA-GFP fusion protein in the control siblings or *cdk1*^−/−^ embryos. PCNA-GFP form puncta during S phase but become diffusive when cells enter G2 phase (Easwaran et al., 2007), which enables us to identify cells in the S phase to G2 phase (Fig. 3D). During S to G2 phase, the retinal nuclei migrate apically. In the *cdk1*^−/−^ retina, 68% of PCNA-GFP puncta-positive nuclei migrate apically, which was comparable to that in the control retina (Fig. 3E). This result confirmed that Cdk1 is not required for IKNM in the zebrafish retina.

### Cdk1 regulates S phase entry of retina cells through Top2a

To further identify the downstream factors of Cdk1 during retinal development, we conducted bioinformatics analysis by searching for the genes included in both the database of Cdk1 substrates and the database of retina expressed genes (Leung et al., 2007). Among 545 Cdk1 substrates, only 77 of them are expressed in zebrafish retina (Fig. 4A). We then manually analyzed the function of the 77 genes, and located *top2a*, which leads to microphthalmia in zebrafish when knocked out (Sapetto-Rebow et al., 2011). Thus, we hypothesized that Top2a may function downstream of Cdk1. Whole mount *in situ hybridization* showed that, similar to *cdk1*, *top2a* was also expressed maternally at 1-cell stage, and then enriched in the retina at 24 hpf and in the CMZ region at 72 hpf (Supplementary Fig. 3). Previous studies show that Top2a regulates cell cycle by regulating DNA architectures (Chen et al., 2015; Deng et al., 2015). To further explore the function of Top2a in Cdk1-dependent cell cycle, we generated a *top2a* mutant by CRISPR/Cas9 technology (Supplementary Fig. 4A). Similar to *cdk1*^−/−^ mutants, *top2a*^−/−^ mutants also showed dorsally curved body (data not shown) and degenerated retinas at 72 hpf (Fig. 4B). Besides, cell proliferation rate was significantly decreased in *Top2a*^−/−^ at 24 hpf (Fig. 4C). Previous studies suggested that Cdk1 phosphorylates Top2a at several Serine residues (Blethrow et al., 2008; Wells. and Hickson, 1995; Xu and Manley, 2007), however, we found only S1213 is conserved cross species (Supplementary Fig. 4B), we tested whether phosphorylation of S1213 was important for Top2a function. Strikingly, overexpression a phosphomimetic form of human TOP2A, TOP2A^S1213D^, but not wild-type TOP2A, rescued microphthalmia of *cdk1*^−/−^ (Fig. 4D). Furthermore, we also observed that the phosphorylation level, but not the transcriptional level, of Top2a in *cdk1*^−/−^ retina 1 was down-regulated (Fig. 4E, Supplementary Fig. 5). The phosphorylation level of Top2a was also significantly down-regulated in *Top2a*^−/−^ (Fig. 4E), showing that the antibody is specific. Thus, our results suggested that Cdk1 functions through Top2a in retinal development.

**Figure 4.**
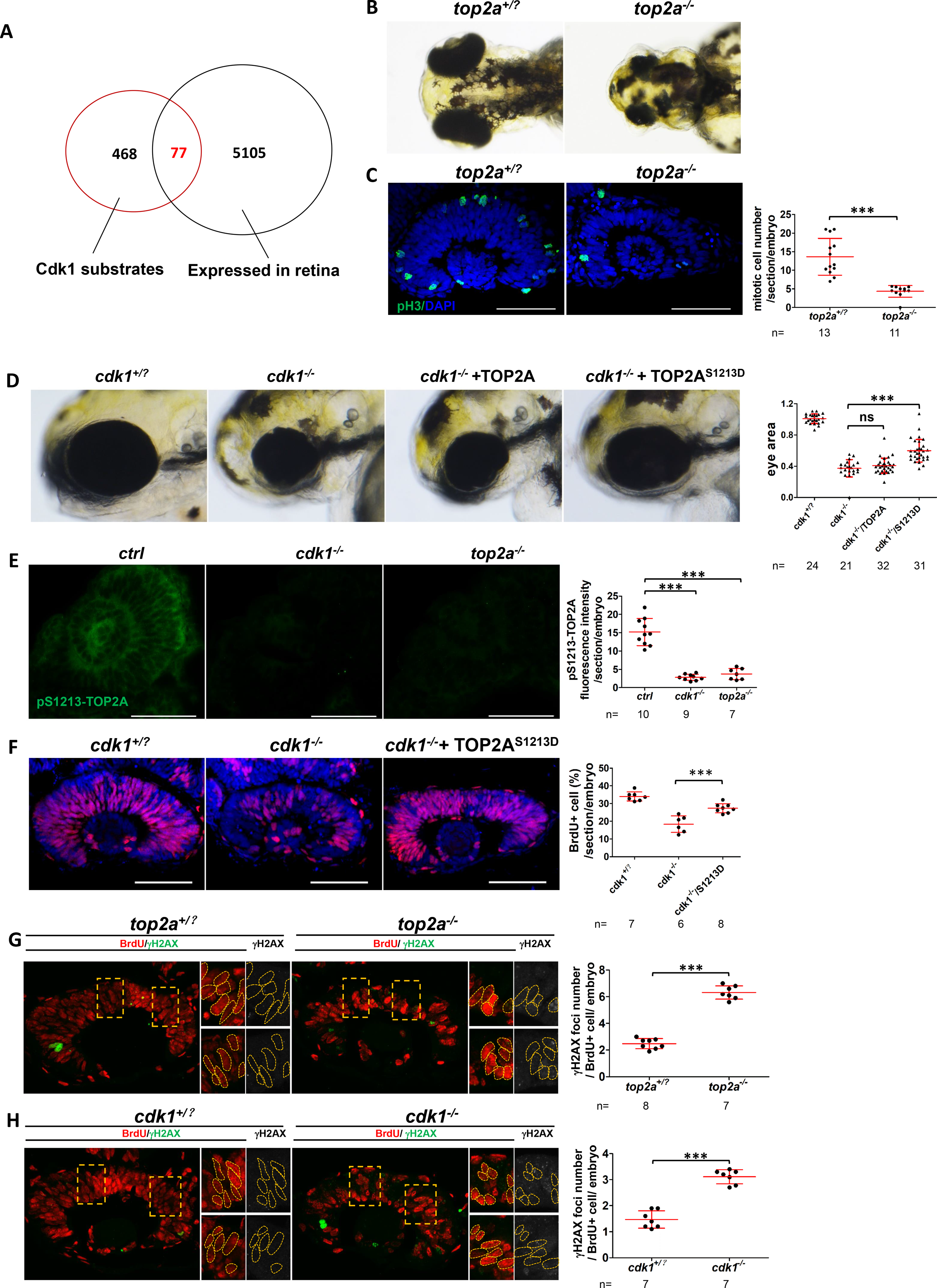
Cdk1 regulates S phase initiation and/or DNA repair by phosphorylating Top2a on S1213 site. (C, E-H) Embryos as indicated sectioned, and retinas were subjected to IF staining with indicated antibodies. (A) Venn Diagram of Cdk1 substrates and retina genes. (B) Images of control siblings and *Top2a*^−/−^ showed the microphthalmia in *top2a^−/−^*. (C) Mitotic cells (pH3) were decreased in *top2a*^−/−^ retinas at 24 hpf. Mitotic cell numbers per section in each embryo were counted and plotted (n≥22 embryos). (D) Microphthalmia of *cdk1*^−/−^ was rescued by phosphorylated (TOP2A^S1213D^) but not wildtype TOP2A. The sizes of eye area were measured and plotted (n≥24 embryos). (E) The level of phosphorylated Top2a on S1213 was decreased in *cdk1*^−/−^ or *Top2a*^−/−^ retinas at 24 hpf. Fluorescence intensity was measured and plotted (n= 7-10 embryos). (F) Cells at S phase (BrdU positive) were decreased in *cdk1*^−/−^ retinas, and rescued by overexpression of TOP2a^S1213D^. Embryos were pulse-treated with BrdU for 1 hour starting from 29hpf. BrdU+ cells percentage was calculated and plotted (n=6-8embryos). (G, H) DNA damage ((H2AX positive puncta) was increase in S-phase cell in *Top2a*^−/−^ (24 hpf) or *cdk1*^−/−^ retinas (30 hpf). γH2AX positive puncta numbers in BrdU+ cells were counted and plotted (> 70 cells from 7 embryos). Error bars represent S.D.; \**P*<0.05, \*\**P*<0.01, \*\*\**P*<0.001.Scale bars, 50 μm.

As cell proliferation was decreased in *cdk1*^−/−^ retina, we asked which phase was affected. Through BrdU pulse-labeling, we detected significantly decreased S phase entry in *cdk1*^−/−^ retinas compared with wild-type retinas. We further asked whether this could also be rescued by overexpression of Top2a^S1213D^. Indeed, DNA incorporation or S phase entry was significantly rescued in Top2a^S1213D^-overexpressed *cdk1*^−/−^ retinas (Fig. 4F). Taken together, Cdk1 functions in S phase entry by phosphorylating Top2a on S1213.

Furthermore, we explored the mechanism of S phase entry that regulated by Cdk1 and Top2a. Top2a regulates cell cycle through relieving the topology stress that occurs during DNA transcription and replication, and Top2a poisons cause DNA double strand break (Berger et al., 1996; Sapetto-Rebow et al., 2011; Wang, 2002). Thus, we asked whether depletion Top2a will cause DNA strand break. By staining an antibody against γH2Ax, a marker for the DNA double strand break (Kuo and Yang, 2008), we discovered that in the *top2a^−/−^*retina, the γH2Ax foci were significantly increased in the S-phase cells (Fig. 4G). Similar to *top2a*^−/−^, the number of γH2Ax foci in the S-phase cells were also significantly increased in the *cdk1* retina (Fig. 4H).Taken together, DNA double strand break was significantly increased in both *cdk1*^−/−^ and *Top2a*^−/−^ retinal cells during S phase entry, suggesting that Cdk1 regulates S phase entry by releasing the topology stress of DNA replication through Top2a.

### Cyclin A2 interacts with Cdk1 to regulate S phase entry through Top2a

During data mining for genes essential for retinal development, we noticed that *ccna2*^−/−^ mutants manifest microphthalmia (Amsterdam et al., 2004), which is similar to *cdk1*^−/−^. Previous studies show that *ccna2* is a downstream 1 gene of G1 to S phase (G1-S) transcriptional network (Bertoli et al., 2013), and functions downstream of Cdk1. Nevertheless, Cyclin A2 can also function as a binding partner to activate Cdk1(Yam et al., 2002). To distinguish these two possible functions of Cyclin A2 in our model, we first checked the expression level of *ccna2* in *cdk1*^−/−^ mutants by RT-PCR or whole mount *in situ* hybridization. From both experiments, *ccna2* expression level remained unchanged in both *cdk1^−/−^* and *Top2a*^−/−^ mutants (Supplementary Fig. 6), suggesting that Cyclin A2 is not regulated by Cdk1 and Top2a at the transcriptional level.

We further examined if Cyclin A2 functions as a partner of Cdk1 to activate Top2a *in vivo*. To test this hypothesis, we generated *ccna2*-mutated embryos by injecting Cas9 protein and *ccna2* gRNA into the zebrafish embryos. We observed the microphthalmia phenotype in the injected embryos, and DNA sequencing suggested that above 70% of the genomic *ccna2* was mutated in phenotypic embryos (Fig. 5, A and B). Besides, similar to *cdk1*^−/−^ and *Top2a*^−/−^ embryos, BrdU incorporations were significantly decreased and γH2Ax-positive cells were increased in the *ccna2*-mutated retina (Fig. 5, C and D). Moreover, consistent with the *cdk1*^−/−^ embryos, phosphorylation of Top2a on S1213 site was significantly reduced in *ccna2*-mutated retina (Fig. 5E). Furthermore, overexpression of constantly phosphorylated Top2a^S1213D^ partially rescued the microphthalmia phenotype of *ccna2*-mutated embryos (Fig. 5F), indicating that Top2a also functions downstream of Cyclin A2.

**Figure 5.**
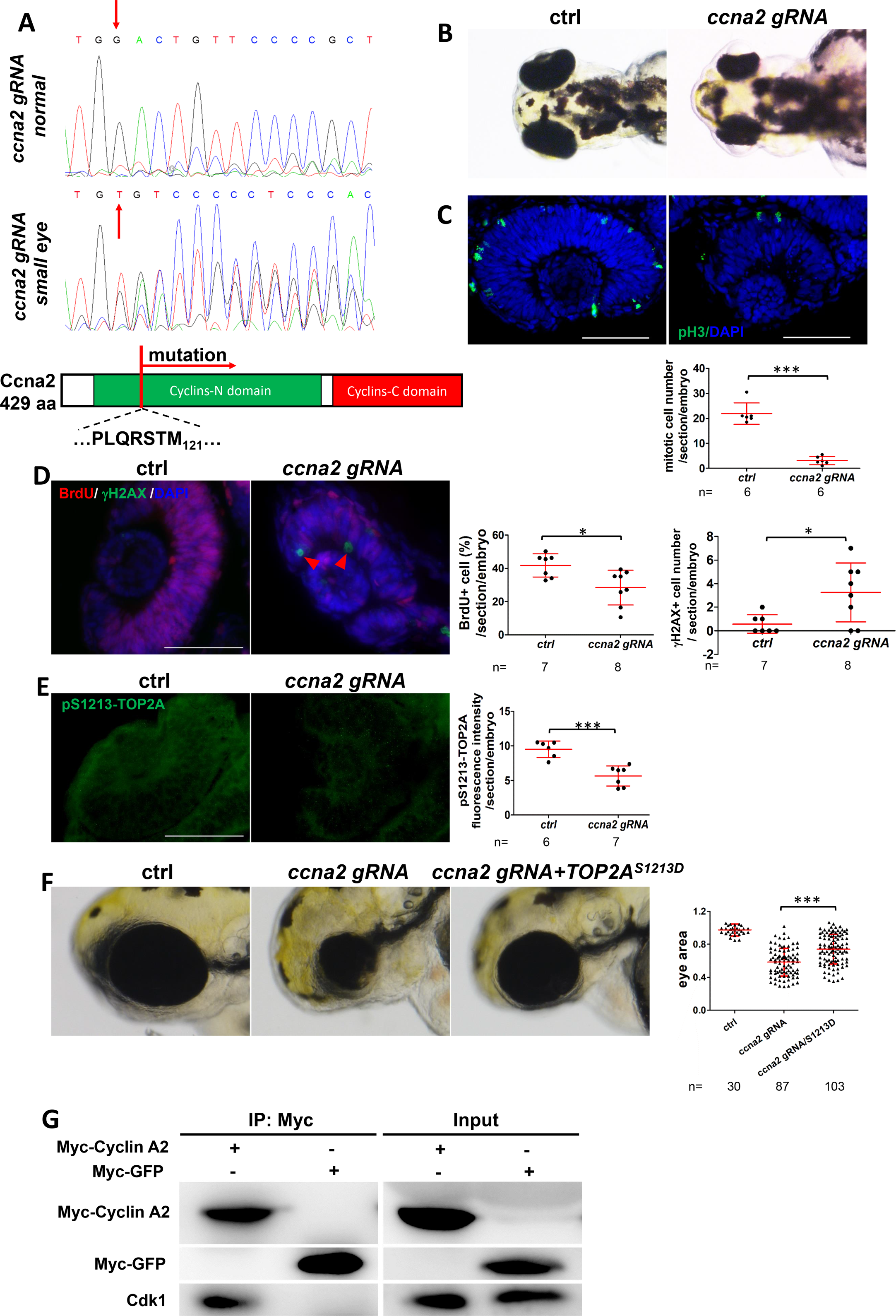
Cyclin A2 functions through Cdk1 and Top2a. (A) Sequence analysis of the *Cas9* mRNA and *ccna2* gRNA injected embryos with or without microphthalmia phenotypes. (B) Light images show microphthalmia in *ccna2* mutants. (C, D, E) Embryos were injected with or without *Cas9* mRNA and *ccna2* gRNA, sectioned, and subjected to IF with indicated antibodies. (C) Mitotic cells (pH3) were decreased in *ccna2* mutant retinas. Mitotic cell number per section in each embryo was counted and plotted (n≥8 embryos). (D) Cells at S phase (BrdU) and with DNA damage response ((H2AX) were increased in *ccna2* mutant retinas. Before collection, embryos were pulse-treated with BrdU for 1 hour start from 23 hpf. BrdU-positive cell percentage and H2AX positive cell numbers per section were counted and plotted (n ≥ 14 section from 7-9 embryos). (E) The level of phosphorylated Top2a on S1213 was decreased in *ccna2* mutant retinas at 24 hpf. Fluorescence intensity was measured and plotted (n≥ sections from 6-7 embryos). (F) Light images of eyes from the control, *ccna2* gRNA-injected, *ccna2* gRNA combined with *TOP2AS1213D* mRNA-injected embryos, as indicated. The sizes of eye area were measured and plotted (n≥0 embryos). (G) Immunoprecipitation result of Myc-Cyclin A2 and Cdk1, in which lysates from the Myc-Cyclin A2 or Myc-GFP overexpressed embryos were immunoprecipitated against Myc antibody and blotted against Cdk1 antibody or Myc antibody. (F) The working model for the role of Cyclin A2, Cdk1 and Top2a in cell cycle regulation. Error bars represent S.D.; \**P*<0.05, \*\**P*<0.01, \*\*\**P*<0.001. Scale bars, 50 μm.

To further verify the interaction between Cyclin A2 and Cdk1, we conducted immunoprecipitation and the results showed that Cyclin A2 physically interact with Cdk1 (Fig. 5G). Taken together, our data unravel the requirements for Cyclin A2-Cdk1-Top2a axis in S phase entry and in preventing DNA double strand break (Fig. 6F).

**Figure 6.**
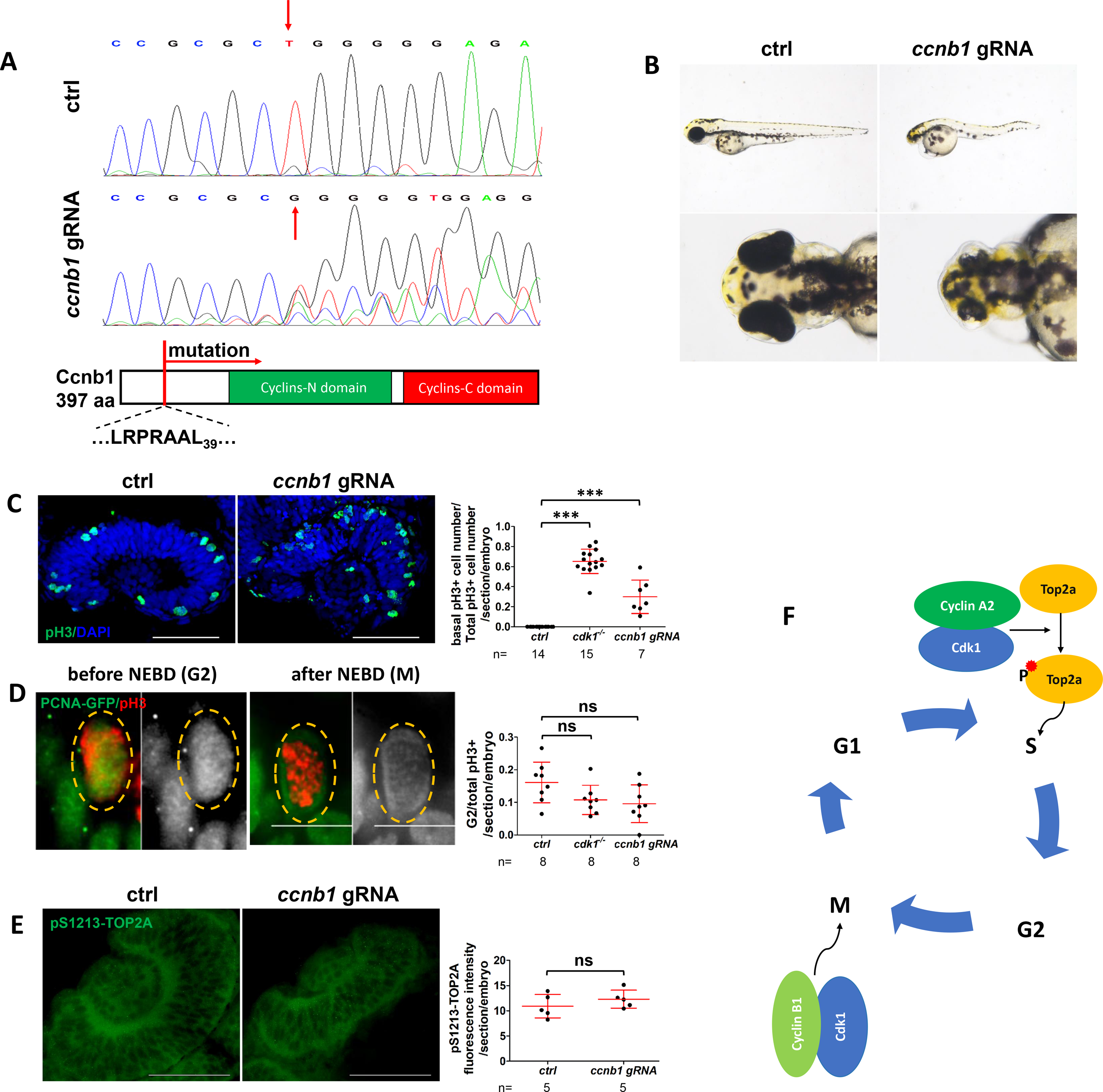
Cyclin B1 is required for M phase progress during retinal development. (A) DNA sequenced of *ccnb1* around the *ccnb1* gRNA target site shows that *ccnb1* gene was mutated in Cas9 protein and *ccnb1* gRNA injected embryos. (B) Microphthalmia in Cas9 protein and *ccnb1* gRNA injected embryos. (C, D, E) Embryos were injected with or without *Cas9* mRNA and *ccnb1* gRNA, sectioned, and subjected to IF with indicated antibodies. (C) Mitotic cells at the basal retina were increased in *ccnb1* mutants. The percentage of mitotic cells at the basal from *cdk1*^−/−^ or *ccnb1* mutant retinas were counted and plotted (n≥7). (D) Representative fluorescent images show cells with PCNA-GFP in the nuclei (before NEBD or at G2 phase) or in the cytoplasm (after NEBD or at M phase). Bar graph shows that the percentage of cells at G2 phase in total pH3 positive cells remained unchanged in *cdk1*^−/−^ or *ccnb1* mutant retinas (n=8). (E) The level of phosphorylated Top2a on S1213 remained unchanged in *ccnb1* mutant retinas at 24 hpf. Fluorescence intensity was measured and plotted (n=5). (F) Schematic model summarizing the results of current study. *in vivo* CyclinA2-Cdk1 complexes phosphorylates Top2a to promote S phase entry, while Cyclin B1-Cdk1 complexes promote M phase process.

### Cyclin B1 functions with Cdk1 to regulate M phase *in vivo*

During data mining, we noticed that mutation in *ccnb1* gene in zebrafish also leads to microphthalmia (Phenotype Annotation (1994-2006)). Cyclin B1, the protein encoded by *Ccnb1*, is the major interacting partner of Cdk1 and catalyzes cell entry into mitosis (Hochegger et al., 2007). Similar to Cdk1-depleted mouse embryos, Cyclin B1-depleted mouse embryos arrest at 4 cell stage, due to G2 arrest accompanied with defects in nuclear envelop break-down (NEBD) (Strauss et al., 2018).

We first validated the previous reported microphthalmia phenotypes observed in Cyclin B1-depleted zebrafish embryos by generating *ccnb1* mutants. By injecting Cas9 protein and *ccnb1* gRNA, we generated *ccnb1*-mutated embryos. DNA sequencing suggested that above 70% of the genomic *ccnb1* was mutated and *ccnb1*-mutated embryos showing microphthalmia as *cdk1*^−/−^ mutants (Fig. 6, A and B).

Next, we analyzed cell cycle progression in *ccnb1*-mutated embryos. Immunostaining with mitotic marker pH3 showed that a significant part of proliferating cells were located at the basal of the retina in ccnb1 mutants (Fig. 5C), which phenocopied *cdk1^−/−^*. As Cyclin B1 is essential for NEBD during early development of mouse embryos (Strauss et al., 2018), we investigated this process in zebrafish. PCNA localized in the nuclear before NEBD and diffused to cytosol after NEBD (Fig. 5D), thus the localization of PCNA can be used as readout for NEBD. In the control retina, 16.1± 6.2% of pH3 positive cells showed nuclear localized PCNA, suggesting that these cells were in the G2 phase (before NEBD), and the rest 73.9% pH3 positive cells showed cytosol localized PCNA, representing cells in the M phase (after NEBD). In both *cdk1*^−/−^ and *ccnb1* mutants, the ratio before NEBD of pH3 positive cells was slightly decreased, but not significant, to that in the control retina, suggesting that NEBD was normal in these two mutants. Taken together, both Cdk1 and Cyclin B1 are required for M phase process of zebrafish retina cells, but not NEBD.

Finally, we asked whether the requirement of Cyclin B1/Cdk1 in M phase is Top2a dependent. By immunostaining, the phosphorylation level of Top2a^S1213^ remained unchanged in *ccnb1*-mutated retinas compared with that in the control retinas. This result suggested that function of Cyclin B1/Cdk1 in M phase was Top2a-independent.

Taken together, our data proved that Cdk1 functioned together with different Cyclins *in vivo* to regulate cell cycle, ie, with Cyclin A2 to regulate S phase entry by phosphorylating Top2a, or with Cyclin B1 to regulate M phase process (Fig. 6F), and its role in cell cycle regulation is essential for vertebrate retinal development.

## Discussion

Cell cycle is an essential cellular process and is tightly regulated by Cyclin-CDK complexes. However, the functions of different Cyclins and CDKs in the different phases of cell cycle are complicated and sometimes controversial (Hochegger et al., 2008). Here, through the zebrafish genetic models, we conducted series of *in vivo* assays, and showed that Cdk1-Cyclin A2 complex is essential for S phase entry through phosphorylation of Top2a. This conclusion is strongly supported by following evidences. First, depletion of Cdk1, Cyclin A2 or Top2a in zebrafish leads to similar phenotypes, including microphthalmia, reduced S phase entry and increased DNA damage in the retina. In consistent, in the mouse model, depletion of Cdk1, Cyclin A2 or Top2a leads to embryonic lethality at very early stage (Akimitsu. et al., 2003; Hochegger et al., 2008; Murphy et al., 1997; Santamaria et al., 2007), but depletion of Cdk2/3/4/6 or Cyclin E1/2, the Cdks or Cyclins considered essential for S phase entry, has little effect on cell cycle progression in mouse (Berthet et al., 2003; Geng et al., 2003; Malumbres et al., 2004; Santamaria et al., 2007; Ye et al., 2001). These findings suggested that the function Cyclin A2-Cdk1-Top2a axis in cell cycle regulation is conserved in vertebrates. The survival of zebrafish mutants during early embryonic development is likely due to the maternal deposits of the gene products in zebrafish embryos, which support early embryonic development. This important feature provides a window to investigate the functions of these genes during late embryonic development. Second, phosphorylation of Top2a on S1213 is reduced in Cdk1- or Cyclin A2-depleted zebrafish retinas, whereas overexpressing TOP2A^S1213D^, a phosphomimetic form of human TOP2A, partially rescues microphthalmia in both *cdk1*^−/−^ and *ccna2*^−/−^. This is consistent with previous studies, which also support the essential role of Cyclin A2, Cdk1, Top2a in S phase (Aleem et al., 2005; Hossain et al., 2004; Kanakkanthara et al., 2016). Third, our and others data have shown that during S phase, Cyclin A2 and Cdk1 physically interact with each other both *in vitro* (Aleem et al., 2005) and *in vivo*, supporting that Cyclin A2 and Cdk1 function together to regulate S phase entry. All these evidences support that Cyclin A2 and Cdk1 functions together to regulate S phase entry by phosphorylation Top2a.

How does Cyclin A2-Cdk1 complex affect S phase entry through Top2a? According to literature and our data, there are at least two possibilities. First, Cyclin A2-Cdk1-Top2a axis regulates S phase entry by preventing DNA damage, which may be caused by mechanical stress possibly accumulated during DNA replication. Supportively, the major role of Top2a is to solving DNA topological problems that are associated with DNA replication (Nitiss, 2009; Wang, 2002). Consistent with this hypothesis, we have shown that DNA damage level is significantly increased in the retina of all the three mutants, *cdk1*^−/−^, *ccna2*^−/−^ and *top2a*^−/−^. Moreover, several studies have shown that Cyclin A2 insufficiency or inhibition of Top2a leads to DNA damage (Deng et al., 2015; Kanakkanthara et al., 2016). Second, as previous studies suggested, Cyclin A2-Cdk1-Top2a axis may regulate the origin firing program, which initiates eukaryotic DNA replication and is a key step in S phase entry (Gaggioli et al., 2013; Kanakkanthara et al., 2016; Katsuno et al., 2009).

Furthermore, Cyclin A2-Cdk1-Top2a axis may also function in the other cell cycle stages besides the S phase entry, for example, chromosome segregation in mitosis. Top2a is well known for its function in chromosome segregation, as depletion or suppression of Top2a leads to lagged or bridged chromosome after mitosis (Akimitsu et al., 2003; Dovey et al., 2009). Similarly, Cyclin A2 deficiency also leads to chromosome segregation defects (Kanakkanthara et al., 2016). However, we did not observe any chromosome segregation defect in *Top2a*^−/−^ mutants, possibly due to maternal deposition of Top2a as suggested in a previous report (Dovey et al., 2009), so we are unable to test this event in our system.

Finally, in our study, we also proved that Cdk1 and Cyclin B1 are required for M phase progression. To our knowledge, for the first time we proved that basally localized pH3 positive cells are M phase arrested cells instead of IKNM-defective cells. These M phase arrested cells exist in Cdk1- or Cyclin B1-depleted retina, but not in Cyclin A2- or Top2a- depleted zebrafish retina, consistent with the model that Cyclin B1-Cdk1 complex function in mitosis (Hochegger et al., 2008).

Taken together, our data revealed that, in an *in vivo* model, Cyclin A2-Cdk1 complex is required for S phase entry by phosphorylating Top2a, and that Cyclin B1-Cdk1 complex is essential for M phase progression.

## Materials and Methods

### Zebrafish husbandry

Wide-type and mutant zebrafish of Tubingen background were maintained according to standard protocol(Westerfield, 2000). Embryos were collected from natural mating and kept at 28.5◻ in E3 solution.

### Mutagenesis by CRISPR/Cas9 system

Generation of zebrafish mutants using the CRISPR/Cas9 system was carried out as previously described(Ota et al., 2014). Cas9 mRNA was in vitro transcribed from a previously reported construct. For *cdk1*, a gRNA was chosen to target the posterior region in the *cdk1* locus. The gRNA targeting sequence is as follows: GGTCTATTTCGGAGTCTCCA. For top2a, a gRNA was chosen to target the middle region in the *top2a* locus. The gRNA targeting sequence is as follows: GAGGTCAATCCCCTGCATGG. Both Cas9 mRNA and gRNA were injected into zebrafish embryos and mutagenesis efficiency was estimated by PCR, amplifying the target sequence from DNA lysate of injected embryos followed by restriction enzyme digestion (Hpy188I for *cdk1*, BslI for *top2a*). For mutagenesis, injected embryos were raised to adulthood and outcrossed to wild-type fish. Positive carrier or founder fish F0 were identified by genotyping the offspring embryos from the outcrosses and F1 were raised. Positive F1 fish were crossed to each other to get homozygous mutant embryos. For*ccna2*, three gRNAs were chosen to target the exon3 in the *ccna2* locus in order to see the phenotype at F0 according to previous publications (Zuo et al., 2017). The gRNA targeting sequences are as follows: GTAAACCTGAAGAAAATGCC, GCTGCTTTTCAGATTCACG, GTCAGCGGGGAACAGTCCA. The mutagenesis efficiency was estimated by sequencing.

### mRNA injection

For mRNA injection, full-length cDNA of *cdk1* was amplified by PCR with EcoRI/XhoI sites from zebrafish cDNA library and inserted into pCS2-linker-gfptogenerate pCS2-*cdk1*-linker-gfp.pCS2-NICD-gfpwas generated from pME-NICD (gift from Dr. Feng Liu) through gateway cloning method. pCS2-Myc-GFP and pCS2-Myc-*ccna2* were also generate through gateway cloning method. The *cdk1-*gfp,NICD-gfp, gfp, Myc-gfp, Myc-*ccna2* and pcna-gfp mRNA were transcribed *in vitro* using the mMESSAGE mMACHINE SP6 kit (Ambion) after linearization of pCS2-*cdk1*-linker-gfp,pCS2-NICD-gfp, pCS2-gfp,pCS2-Myc-gfp,pCS2-Myc-*ccna2* and pCS2-pcna-gfp (from Dr. Jie He) constructs respectively with NotI (Takara). pcDNA3.1-TOP2A (WT) and pcDNA3.1-TOP2A (S1213D) were the gifts from Dr. Santocanale Corrado. pcDNA3.1-TOP2A (S1354D) was generated using pcDNA3.1-TOP2A (WT) as template by Fast Mutagenesis System (transgene). TOP2A (WT)/TOP2A (S1213D)/TOP2A (S1354D) mRNA were transcribed in vitro after linearization of constructs with BamHI (Takara) using the mMESSAGEmMACHINET7 kit (Ambion).

### In Situ Hybridization

Whole-mount in situ hybridization was performed as previously described. Full length of *ccna1/ ccna2/ ccne1* and the posterior 1738bp of *top2a* were amplified by PCR from zebrafish cDNA library with a T7 sequence on the reverse primer. Antisense probes for *ccna1/ ccna2/ ccne1/ top2a* were made using the PCR products as templates for RNA synthesis with T7 RNA polymerases (Takara). The stained embryos were dehydrated in glycerol and photographed with a Nikon SMZ1500 stereomicroscope (Nikon, Tokyo, Japan).

### Histology

To determine the cellular pattern of the *cdk1* mutant retina, we fixed embryos in 4% formaldehyde in 1×PBST at RT for 6 hours then dehydrated and embedded the tissues in OCT at −80◻ overnight. Embryos were sectioned at 8μm thickness using a Leica cryostat. The tissues were stained with hematoxylin for 30s, the rest steps are according to previous publications (Ellis and Yin, 2017).

### Antibody staining

Embryos were fixed with 4% formaldehyde solution for 2 hours at RT. Samples were blocked with blocking buffer (10% serum in PBST) and incubated with primary antibodies in blocking buffer. Primary antibodies were recognized by secondary antibodies conjugated with Alexa Flour 488 or Alexa Flour 594. The images were taken with a Nikon eclipse 80i fluorescent microscope.

Primary antibodies were diluted in blocking buffer at following concentrations: anti-zpr1 (1:200) and anti-zpr3 (1:500) were gifts from Dr. Chengtian Zhao, anti-pH3 (1:1000, Millipore),anti-GFP (1:500, Roche), anti-β-tubulin (1:300, Sigma-Alorich), anti-BrdU (1:500, Sigma-Aldrich), anti-γH2Ax (1:300, CST), anti-γ-tubulin (1:200, Invitrogen), anti-aPKC (1:200, Santa Cruz), anti-phos-S1213-TOP2A (1:300, Invitrogen). The secondary antibodies (Invitrogen) were used at 1:500 dilutions.

### TUNEL assay

For TUNEL assay, embryos were fixed with 4% formaldehyde solution for 2 hours at RT, then dehydrated and embedded the tissues in OCT at −80◻ overnight. Embryos were sectioned at 10μm thickness using a Leica cryostat. The tissues were stained with an In Situ Cell Death Detection Fluorescein kit (Roche) according to the manufacturer’s instructions.

### BrdU pulse-labeling

Embryos were incubated for 20 min in BrdU (10◻mM, in embryo medium with 15% DMSO) at 4 ◻. For *cdk1* siblings and mutants, embryos were incubated with BrdU at 29◻hpf. For *top2a* siblings and mutants and *ccna2* gRNA injected embryos, embryos were incubated with BrdU at 23◻hpf. After BrdU treatment, embryos were cultured for extra 1hour in embryo medium at 28.5◻ and fixed with 4% formaldehyde. Embryos were then cryosectioned and carried out for immunofluorescent staining.

### Quantitative real-time PCR

Total cellular RNA was extracted using the TRIzol reagent (Takara). Complementary DNA was synthesized with 1μgtotal RNA using a PrimeScript™ RT-PCR Kit (Takara). The real-time PCR with the mixture reagent SYBR Green (Thermo Fisher Scientific) was carried out on a real-time detection system (ABI7500). Gene specific primers were used. Cycling conditions are 50◻ for 2min, 95◻ for 2min, followed by 40 cycles of 95◻ for 15s and 60◻ for 1min. For quantification, the target genes were normalized against the amount of beta-actin. Primers for real-time PCR is as follows: *ccna1*: forward (5’→3’): ACTCGACGATGCTGTTCAAGATA, reverse (5’→3’): GAAAGAAAGCGGTCCAGGTAAT; *ccna2*: forward (5’→ 3’): GGACTGGTTGGT GGAAGTGG, reverse (5’ → 3’:ACTGATTGATGGTGGGAGCG; *ccne1*: forward (5’ → 3’):AGTTTGCTTATGTTACTGATGGG G, reverse (5’ → 3’):GAGAAGAAA AGTGGAAGAGTGCTG; *top2a*: forward (5’ → 3’):CCCCAAGAGAAAAGAGAG AACG, reverse (5’ → 3’):CAACAGGTGGAGGAGGAGAGTC.

### Western Blot and Co-Immunoprecipitation

The fish embryos were lysed in a buffer containing 4% SDS, 10mM Na phosphate utter (pH 7.0), 40% glycerol, 0.2% bromophenol blue, 0.2 M DTT and 5% mercaptoethanol, then boiled at 95◻ for 10min. Embryos of the total protein lysates were used for western blot analysis using anti-Cdk1 antibody (1:300, Santa Cruz) and anti-beta-Actin antibody (1:1000, Santa Cruz). The second antibody is goat anti-mouse HRP (1:2000, Abgent). All of the antibodies were diluted in 3% BSA/PBST.

For Co-immunoprecipitation, the fish embryos injected with Myc-ccna2 mRNA or Myc-gfp mRNA were collected and lysed at bud stage. The lysis solutions were incubated with anti-Myc antibody (2μl in each lysis) at 4 □overnight then incubated with protein A beads at 4 ◻ for an hour. The beads were washed and the Myc-Cyclin A2, Myc-GFP and endogenous Cdk1 proteins were detected by immunoblotting.

### In vivo time-lapse imaging

Embryos were treated with 0.04% Tricainemesylate (MS-222) in 0.003% PTU and embedded in 1% low melt agarose in E3 medium on a glass-bottom culture dish. For GFP labeled cells, the experiment was carried out on Olympus FV1200 inverted confocal microscope with 30× (oil, NA= 1.05) objectives. For PCNA-GFP labeled cells, the experiment was performed with 60× (NA=1.20) objectives. A 28◻ heating chamber was used throughout the entire time-lapse. Optional section of Z-stacks was collected at 2μm every 20min for 6 hours. Time-lapse experiments were started with embryos at 28 hpf.

### Statistical analysis

Data are presented as mean ± SD. Statistical differences between two sets of data were analyzed using a two-tailed paired student’s t-test and a value of *p* < 0.05 was considered significant.

## Acknowledgements

We thank Dr. Lu Zhao (Zhejiang University), Dr. Qiang Chen (Wuhan University) for critical reading the manuscript, the members of the Cao lab and He lab for helpful discussions. This project was supported by grants from National Key Research and Development Program of China (2017YFA0104600), National Natural Science Foundation of China (31771597 and 31571515),National Basic Research Program of China (973 Program 2012CB966601and 2013CB945300), Ministry of Science and Technology of China (2011DFB30010).

