## supplemental data for "Cyclin A2/Cdk1 is Essential for the *in vivo* S Phase Entry by Phosphorylating Top2a"

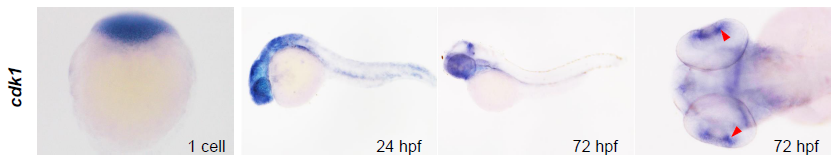

**Supplemental Figure 1.***Cdk1* is maternally supplied and zygotically expressed at retinal CMZ region. The expression patterns of *cdk1* were revealed by *in situ hybridization* in wild-type embryos at indicated stages. CMZ, ciliary marginal zone.

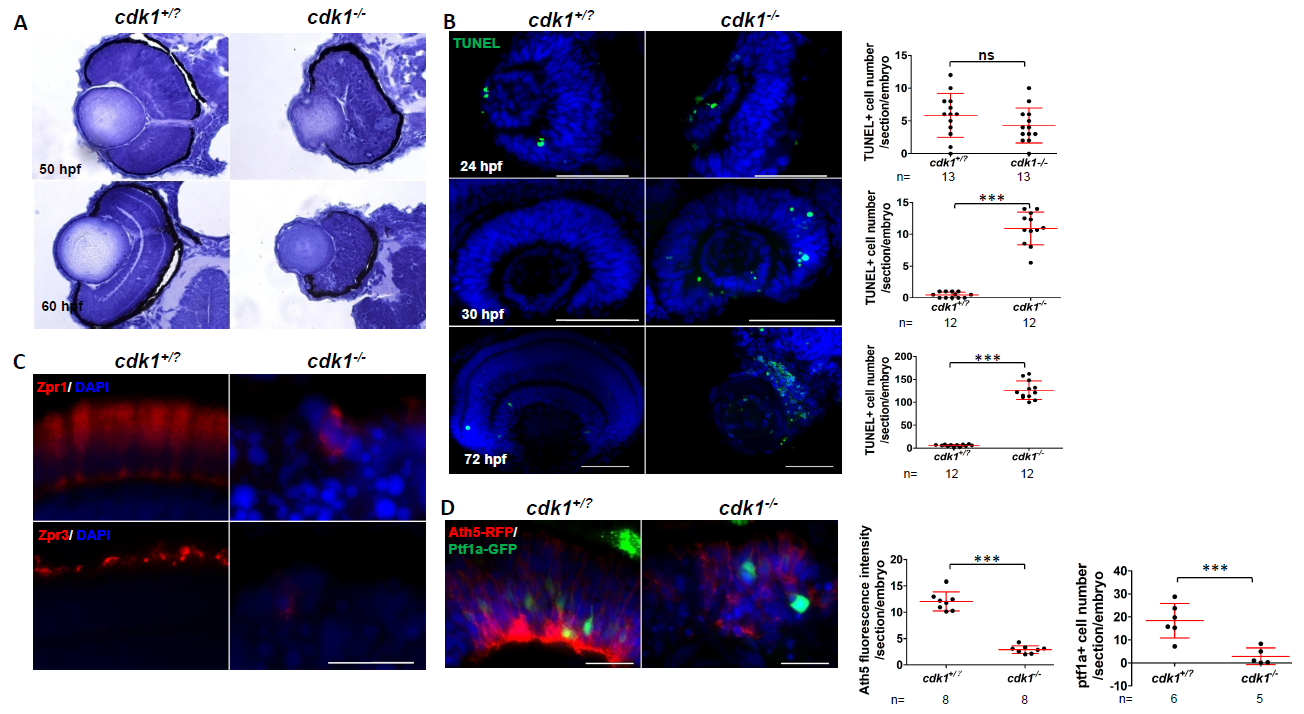

**Supplemental Figure 2**. Retinal development defects in *cdk1^-/-^*. (A-C) Embryos were cross-sectioned and the retinas were stained with hematoxylin, or subjected to IF microscopy with indicated antibodies. (A) Retinal lamination is disrupted in the *cdk1^-/-^*embryos. Embryos were cross-sectioned and the retinas were stained with hematoxylin. (B) Cell apoptosis is increased in *cdk1^-/-^* retinas at 30 hpf or 72 hpf, but not 24 hpf. Apoptotic cell numbers per section were counted and plotted (n≥12 embryos). (C) Photoreceptor cells (Zpr1 for rod cells and Zpr3 for cone cells) were lost in *cdk1^-/-^* retinas at 3 dpf. (D) Ath5-RFP positive and Ptf1a-GFP positive cells were decreased in *cdk1^-/-^* retinas. Fluorescence intensity or positive cells per section were measured and plotted (n≥5 embryos). Error bars represent S.D.; **P*<0.05, ***P*<0.01, ****P*<0.001.Scale bars, 20 μm in C and D; 50μm in B.

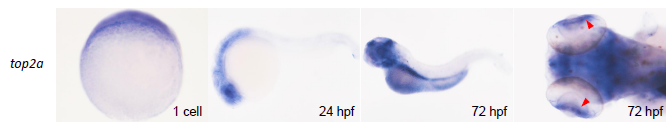

**Supplemental Figure 3.** *top2a* is maternally supplied and zygotically expressed at retinal CMZ region. The expression patterns of *top2a* were revealed by *in situ hybridization* in wild-type embryos at indicated stages. CMZ, ciliary marginal zone.

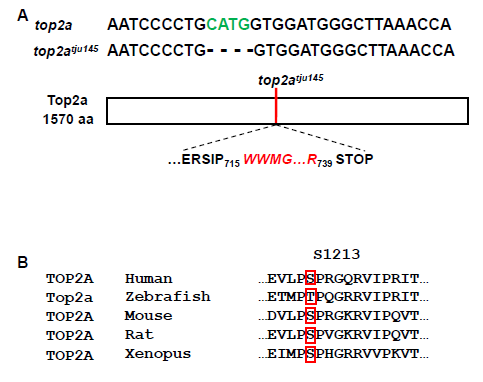

**Supplemental Figure 4**. null mutant of *top2a*. (A) DNA sequences of the wild-type and the mutant *top2a* alleles. Schematic of Top2a protein indicates position of the *top2a^tju145^* frameshift, which leads to premature stop codon. (B) Sequence comparison shows the conservation on S1213 of Top2a across species.

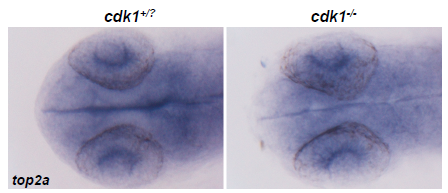

**Supplemental Figure 5**. The expression level of *top2a* remains unchanged in *cdk1^-/-^* retinas. Embryos were subjected to whole mount *in situ* hybridization.

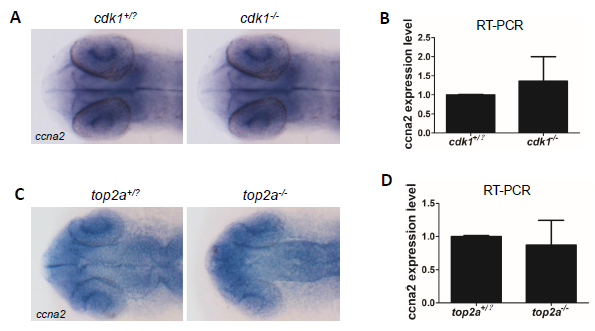

**Supplemental Figure 6**. The expression level of *ccna2* remains unchanged in *cdk1^-/-^* or *top2a^-/-^*. (A-D) Both *in situ hybridization* (A and C) and RT-PCR (B and D) revealed that the expression level of *ccna2* remained unchanged in *cdk1^-/-^* or *top2a^-/-^*.
